# DNA methylation as a marker for prenatal smoke exposure in adults

**DOI:** 10.1101/121558

**Authors:** R.C. Richmond, M. Suderman, R. Langdon, C.L. Relton, Smith G. Davey

## Abstract

Prenatal cigarette smoke is an environmental stressor that has a profound effect on DNA methylation in the exposed offspring. We have previously shown that some of these effects persist throughout childhood and into adolescence. Of interest is whether these signals persist into adulthood.

We conducted an analysis to investigate associations between reported maternal smoking in pregnancy and DNA methylation in peripheral blood of women in the Avon Longitudinal Study of Parents and Children (ALSPAC) (n=754; mean age 30 years). We observed associations at 15 CpG sites in 11 gene regions, *MYO1G, FRMD4A, CYP1A1, CNTNAP2, ARL4C, AHRR, TIFAB, MDM4, AX748264, DRD1, FTO* (FDR < 5%). All but two of these CpG sites have previously been identified in relation to prenatal smoke exposure in the offspring at birth and the majority showed persistent hypermethylation among the offspring of smokers.

We confirmed that most of these associations were not driven by own smoking and that they were still present 18 years later (N = 656; mean age 48 years). In addition, we replicated findings of a persistent methylation signal related to prenatal smoke exposure in peripheral blood among men in the ALSPAC cohort (N = 230; mean age 53 years). For both participant groups, there was a strong signal of association above that expected by chance at CpG sites previously associated with prenatal smoke exposure in newborns (Wilcoxon rank sum p-value < 2.2 × 10^−4^). Furthermore, we found that a prenatal smoking score, derived by combining methylation values at these CpG sites, could predict whether the mothers of the ALSPAC women smoked during pregnancy with an AUC 0.69 (95% 0.67, 0.73).

## Introduction

Cigarette smoke exposure during pregnancy is an environmental stressor that has a profound effect on DNA methylation in the exposed offspring (1). Previous work has determined genome-wide changes in DNA methylation in response to smoke exposure in-utero (2), with a recent epigenome-wide association study (EWAS) meta-analysis of methylation in newborn cord blood identifying over 6,000 differentially methylated CpG sites (of which 568 CpG sites met the strict Bonferroni threshold for statistical significance) (3).

It is of interest to investigate the persistence of methylation marks into later life as this presents the opportunity to use methylation as an archive of historical exposure, particularly if methylation patterns can be robustly modelled over time (4, 5). In addition, persistent changes in DNA methylation might mediate at least some of the associations between smoke exposure in pregnancy and later life health outcomes (6).

Several studies have identified prenatal smoke exposure associated changes in methylation in childhood and adolescence in global methylation (7), candidate gene (7-9) and epigenome-wide association studies (EWAS) (3, 4, 10-13). However, few studies have investigated the persistence of methylation change into adulthood (14-16). Two such studies have been conducted in the multi-ethnic New York City birth cohort, where prospectively assessed maternal smoking during pregnancy was found to be positively associated with global methylation in leukocytes of individuals at age 43 years, assessed using a methyl acceptance assay (14), and inversely associated with levels of *Sat2* methylation (15), which remained even after adjustment for adult smoking status of the offspring. However, a more recent EWAS found no single sites associated with prenatal smoke exposure in lymphoblastoid cell lines (LCL) of male offspring at age 45 years, although one differentially methylated region (DMR) was found to be weakly associated in whole blood (16). All of these previous studies were limited by low power due to small sample sizes and a more comprehensive assessment of the long-term impact of prenatal tobacco smoke exposure on genome-wide methylation is warranted.

In this study, we aimed to assess the long-term impact of prenatal tobacco smoke on DNA methylation in the context of the Avon Longitudinal Study of Parents and Children (ALSPAC) (17, 18), a prospective birth cohort with data on reported maternal smoke exposure in pregnancy and genome-wide DNA methylation levels of offspring in adulthood. We first conducted an analysis to investigate associations between reported prenatal smoke exposure and DNA methylation in peripheral blood among women in ALSPAC. We next attempted to replicate prenatal smoking-associated DNA methylation differences in peripheral blood of the women 18 years later and in men from the same study (the partners of these women). Finally, we aimed to assess the extent to which a prenatal smoking score, based on methylation at CpG sites previously shown to be associated with prenatal smoke exposure, could predict whether the mothers of the ALSPAC women smoked during pregnancy.

## Methods

### Cohort and selection of participants

ALSPAC is a large, prospective cohort study based in the South West of England. 14,541 pregnant women resident in Avon, UK with expected dates of delivery 1st April 1991 to 31st December 1992 were recruited and detailed information has been collected on these women and their offspring at regular intervals (17, 18). The study website contains details of all the data that are available through a fully searchable data dictionary (http://www.bris.ac.uk/alspac/researchers/data-access/data-dictionary/). Written informed consent has been obtained for all ALSPAC participants. Ethical approval for the study was obtained from the ALSPAC Ethics and Law Committee and the Local Research Ethics Committees.

We examined offspring DNA methylation in relation to reported maternal smoking during pregnancy in ALSPAC using methylation data from the Illumina Infinium HumanMethylation450 (HM450) BeadChip assay. We used data from women enrolled ALSPAC (n=754) in the main analysis and looked to replicate findings in the same women approximately 18 years later (n=656), and in men enrolled in ALSPAC (n=230).

### Prenatal exposure variables

After recruitment of pregnant women into the ALSPAC study, information was collected on both the women and their partners, including details of their mothers’ smoking behaviour. If the men and women reported that their mothers had smoked, they were asked whether their mothers had smoked when they were pregnant with them, and if so, were given the responses yes/no/don’t know from which to select. These data were analysed assuming that, for all those who said don’t know, their mothers’ did smoke during pregnancy, as has been done previously (19).

Information on the women’s own smoking status was obtained from a questionnaire administered at 18 weeks’ gestation. Women were asked whether they had smoked regularly pre-pregnancy, from which a dichotomous variable for any tobacco smoking before pregnancy was derived. Information on the women’s smoking status was also gathered in a questionnaire administered approximately 18 years later. Furthermore, information on the partner’s smoking status was obtained from a questionnaire administered approximately 21 years after the pregnancy. At these later time points, the men and women were asked whether they currently smoked or whether they had smoked every day when a smoker in the past. From these data, a dichotomous variable for any previous tobacco smoking was derived.

### DNA methylation assessment

We examined offspring DNA methylation in peripheral blood (whole blood, white cells, peripheral blood lymphocytes [PBLs]) in ALSPAC men and women.

As part of the ARIES project (20), the Illumina Infinium HumanMethylation450K (HM450) BeadChip (21) has been used to generate epigenetic data on 1,018 mother-offspring pairs in the ALSPAC cohort. A web portal has been constructed to allow openly accessible browsing of aggregate ARIES DNA methylation data (ARIES-Explorer) (http://www.ariesepigenomics.org.uk/).

The ARIES participants were selected based on the availability of DNA samples at two time points for the women (antenatal and at follow-up when the offspring were adolescents) and three time points for their offspring (neonatal, childhood [mean age 7.5 years], and adolescence [mean age 17.1 years]). Furthermore, additional 450K data has been generated on ALSPAC men (the partners of the women enrolled in ARIES) (n=312).

We examined DNA methylation in relation to prenatal smoke exposure using methylation data from the Infinium HM450 BeadChip. For the samples, the methylation level at each CpG site was calculated as a β-value, which is the ratio of the methylated probe intensity and the overall intensity and ranges from 0 (no cytosine methylation) to 1 (complete cytosine methylation). All analyses of DNA methylation used these β-values.

Cord blood and peripheral blood samples (whole blood, buffy coats, or blood spots) were collected according to standard procedures, and the DNA methylation wet-laboratory and pre-processing analyses were performed as part of the ARIES project, as previously described (20). In brief, samples from all time points in ARIES were distributed across slides using a semi-random approach to minimize the possibility of confounding by batch effects. Samples failing quality control (average probe *P* value ≥0.01, those with sex or genotype mismatches) were excluded from further analysis and scheduled for repeat assay, and probes that contained <95% of signals detectable above background signal (detection *P* value <0.01) were excluded from analysis. Methylation data were pre-processed using R software, with background correction and subset quantile normalization performed using the pipeline described by Touleimat and Tost (22).

### Covariates

Maternal age at birth and head of household social class were included as covariates in these analyses as they were found to be most strongly associated with smoking status during pregnancy in a previous study (4). Information on the ALSPAC parents’ mothers’ age at delivery was recorded in a questionnaire administered at 18 weeks’ gestation. Information on the ALSPAC parents’ fathers’ social class was also recorded in the same questionnaire and was classified into non-manual or manual work. We also adjusted for current age at blood draw for methylation profiling among the participants. 10 surrogate variables (SVs) were generated using the “SVA” package in R and included in models to adjust for technical batch and cell-type mixture (23). This was done given the heterogeneity in cell types between the different samples. We also included smoking behaviour of the adult offspring as a further covariate in sensitivity analyses to consider the potential influence of own smoking which might explain the persistence in methylation signatures associated with intrauterine exposure.

### Statistical analysis

We first performed an epigenome-wide association analysis in our largest sample, the ALSPAC women with methylation measured in peripheral blood taken at the time of enrolment in the study (n=754). CpG level methylation (untransformed beta-values) was regressed against prenatal smoke exposure (any maternal smoking during pregnancy) with adjustment for covariates (maternal age, parental social class, offspring age, top 10 SVs). We then assessed whether associations were robust to adjustment for own smoking status and further performed replication analyses for the top CpG sites (false discovery rate [FDR] < 0.05) in the other samples, as well as assessing the association between prenatal smoke exposure and methylation at these sites in cord blood in the ALSPAC cohort which has been investigated previously (3, 4).

We also investigated the association between prenatal exposure to smoking and DNA methylation for the 568 CpG sites previously found to be robustly associated with prenatal smoke in a cord blood meta-analysis (n=6,685) (3) in each of the adult cohorts. We assessed the degree of inflation of association signal (lambda value) for these CpG sites compared with that seen genome wide across the samples, and performed a Wilcoxon rank sum test to assess enrichment.

Furthermore, we generated a DNA methylation score (24) for prenatal smoking based on these 568 CpG sites and compared its ability to predict whether the mothers of the ALSPAC adults smoked during pregnancy with a score based on 19 CpG sites which reached Bonferroni significance in a EWAS of prenatal smoking conducted in peripheral blood of older children (n=3,187) (3) and a score for own smoking, consisting of 2,623 CpG sites which reached Bonferroni significance in the largest EWAS of own smoking to date (n=15,907) (25). To derive the scores, we used methylation data for the largest group of participants: the women in ALSPAC at the time of enrolment (n=754). For each individual, a weighted score was obtained by multiplying the methylation value at a given CpG by the effect size from the previous EWAS, and then summing these values:

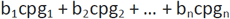
where “cpg” is the normalized methylation value in the ARIES women and “b” is the effect size from the aforementioned smoking EWAS (3, 25). We generated receiver operating characteristic (ROC) curves for the prenatal and own smoking methylation scores and calculated the area under the curve (AUC) in order to assess the performance of these predictors.

We also investigated the extent to which the prenatal smoking scores could predict maternal smoking in pregnancy independent of own smoking status. This was done by comparing methylation scores between four different groups of participants, determined based on their own smoking status and that of their mothers during pregnancy: non-smokers whose mothers never smoked in pregnancy, smokers whose mothers never smoked in pregnancy, non-smokers whose mothers smoked in pregnancy and smokers whose mothers smoked in pregnancy.

Analysis was performed using Stata (version 14) and R (version 3.3.1).

## Results

The cohort-specific summary statistics for this analysis are presented in **Table 1**. The ALSPAC women in our main analysis had a mean age 30 years, while their mean age at follow-up was 48 years and the ALSPAC men had a mean age of 53 years. Maternal age at birth was similar between the ALSPAC men and women, as were parental social class and rates of prenatal smoke exposure. Rates of own smoking varied quite substantially, from 16.9% in the ALSPAC women at the first time point to 33.8% in the ALSPAC men.

**Table 1:** Descriptive characteristics of participant groups in this study

|  | ALSPAC adult females<br>(time point 1) | ALSPAC adult<br>females (time point<br>2) | ALSPAC adult males |
| --- | --- | --- | --- |
| N | 754 | 656 | 230 |
| Maternal age at birth<br>(years) (SD) | 27.6 (5.7) | 27.8 (5.7) | 28.2 (5.7) |
| Social class (manual)<br>(N, %) | 347 (46.0) | 301 (45.9) | 92 (40.0) |
| Prenatal smoke<br>exposure (yes) (N, %) | 216 (28.7) | 179 (27.3) | 73 (31.7) |
| Age at follow-up<br>(years) (SD) | 30.3 (4.3) | 48.1 (4.2) | 53.4 (5.1) |
| Sample type(s) | Whole blood/white cells | White cells/PBLs | White cells/PBLs |
| N | 752 | 498 | 222 |
| Own smoking (N, %) | 127 (16.9) | 144 (28.9) | 75 (33.8) |

We observed associations between 15 CpG sites and prenatal smoking exposure in women at age 30 (FDR < 5% for ∼485,000 tests; **Table 2**). These sites were located in 11 gene regions and all but two have been previously identified in EWAS for maternal smoking at birth (3) and into childhood/adolescence (3, 4, 9-12) and all agree on direction of effect (**Supplementary Table 1**). For example, CpG sites located at *FRMD4A* (cg11813497), *CYP1A1* (cg05549655)*, CNTNAP2* (cg25949550)*, AHRR* (cg17924476), *TIFAB* (cg11429111), *AX748264* (cg11641006) and *FTO* (cg26681628) have all been implicated in the largest cord blood EWAS of prenatal smoking to date (3), and those at *FRMD4A (10, 12), CYP1A1* (4, 9, 11), *CNTNAP2* (4, 9, 11, 12) and *AHRR* (4, 9, 11) have also found to be differentially methylated among older children.

**Table 2:** DNA methylation changes associated with prenatal smoke exposure in ALSPAC women

| CpG site | Chromosome | Gene region | Position | Basic model (N=754) |  |  |  | Adjusted model* (N=752) |  |  |  |
| --- | --- | --- | --- | --- | --- | --- | --- | --- | --- | --- | --- |
|  |  |  |  | Effect size | Standard error | P-value | FDR | Effect size | Standard error | P-value | FDR |
| cg22132788 | 7 | *MYO1G* | 45002486 | 0.062 | 0.008 | 1.70E-13 | 8.26E-08 | 0.058 | 0.008 | 9.77E-13 | 4.75E-07 |
| cg12803068 | 7 | *MYO1G* | 45002919 | 0.111 | 0.015 | 9.82E-13 | 2.38E-07 | 0.097 | 0.014 | 1.71E-11 | 4.16E-06 |
| cg11813497 | 10 | *FRMD4A* | 14372879 | 0.038 | 0.006 | 1.25E-10 | 2.02E-05 | 0.036 | 0.006 | 4.85E-10 | 7.85E-05 |
| cg04180046 | 7 | *MYO1G* | 45002736 | 0.041 | 0.006 | 2.53E-10 | 3.07E-05 | 0.037 | 0.006 | 3.43E-09 | 3.33E-04 |
| cg05549655 | 15 | *CYP1A1* | 75019143 | 0.007 | 0.001 | 1.21E-08 | 0.001 | 0.007 | 0.001 | 8.82E-10 | 1.07E-04 |
| cg25949550 | 7 | *CNTNAP2* | 145814306 | -0.005 | 0.001 | 1.48E-08 | 0.001 | -0.004 | 0.001 | 2.41E-07 | 0.015 |
| cg19089201 | 7 | *MYO1G* | 45002287 | 0.038 | 0.007 | 1.75E-08 | 0.001 | 0.034 | 0.007 | 1.88E-07 | 0.013 |
| cg05204104* | 2 | *ARL4C* | 235403141 | 0.023 | 0.004 | 2.30E-08 | 0.001 | 0.023 | 0.004 | 4.12E-08 | 0.003 |
| cg17924476 | 5 | *AHRR* | 323794 | 0.045 | 0.008 | 6.24E-08 | 0.003 | 0.040 | 0.008 | 5.33E-07 | 0.029 |
| cg11429111 | 5 | *TIFAB* | 134813329 | 0.022 | 0.004 | 2.62E-07 | 0.013 | 0.019 | 0.004 | 2.90E-06 | 0.108 |
| cg08241939* | 1 | *MDM4* | 204700816 | 0.022 | 0.004 | 5.04E-07 | 0.022 | 0.019 | 0.004 | 3.40E-06 | 0.110 |
| cg11641006 | 2 | *AX748264* | 235213874 | 0.040 | 0.008 | 5.62E-07 | 0.023 | 0.029 | 0.006 | 1.27E-06 | 0.051 |
| cg15016771* | 2 | *ARL4C* | 235403218 | 0.008 | 0.002 | 1.01E-06 | 0.036 | 0.007 | 0.002 | 3.35E-06 | 0.110 |
| cg22807681* | 5 | *DRD1* | 174622933 | 0.023 | 0.005 | 1.04E-06 | 0.036 | 0.023 | 0.005 | 1.02E-06 | 0.045 |
| cg26681628 | 16 | *FTO* | 54210550 | 0.037 | 0.008 | 1.27E-06 | 0.041 | 0.029 | 0.006 | 8.94E-06 | 0.193 |

\*Model includes adjustment for own smoking.

Associations at CpG sites located near *ARL4C* (cg05204104 and cg15016771), *MDM4* (cg08241939) and *DRD1* (cg22807681) appear to be novel, although the sites at *ARL4C* were present in the extended list of FDR significant sites in the previous cord blood EWAS (3). In contrast to findings in cord blood, where smoking during pregnancy has been associated approximately equally with hyper and hypomethylation (3), the majority (14 out of 15 CpGs) showed long-term hypermethylation in this analysis.

While in most EWAS for cord blood methylation *AHRR* (cg05575921) is the CpG site most consistently associated with prenatal smoke exposure (3, 4, 26, 27), this site did not survive adjustment for multiple tests in our persistence analysis (P = 0.0005, FDR > 5%). Rather, associations at four sites at *MYO1G* (cg22132788, cg12803068, cg04180046 and cg19089201) did survive multiple testing adjustment, with the *MYO1G* site cg22132788 having the strongest association (P = 1.7×10^−13^), which is consistent with findings in children and adolescents (4, 11, 12).

One concern related to the identification of these signals is that they might reflect non-specific smoke exposure of the offspring over the life course rather than a “critical period” effect of smoke exposure in utero (28). In particular, 11 out of the 15 CpGs have been identified in relation to current versus never smoking status at FDR<0.05 in a recent EWAS of own smoking (25) (**Supplementary Table 2**). To account for this, we examined whether past cigarette smoking by the adult themselves influenced these associations by including own smoking history (yes/no) as a covariate in the model. Adjustment for own smoking attenuated associations at five CpG sites (*TIFAB* (cg11429111)*, MDM4* (cg08241939)*, AX748264* (cg11641006), *ARL4C* (cg15016771) and *FTO* (cg26681628)) so as to no longer reach the FDR cut-off for significance (**Table 2**, **Figure 1**), although on the whole the magnitude of effect was only slightly reduced with this adjustment. Furthermore, while methylation of CpG sites at *MYO1G, CNTNAP2* and *AHRR* have been consistently identified in relation to own smoking, no such associations have been found with CpG sites at *FDRM4A, CYP1A1, MDM4* or *DRD1* in the largest EWAS of own smoking to date (29).

**Figure 1:**
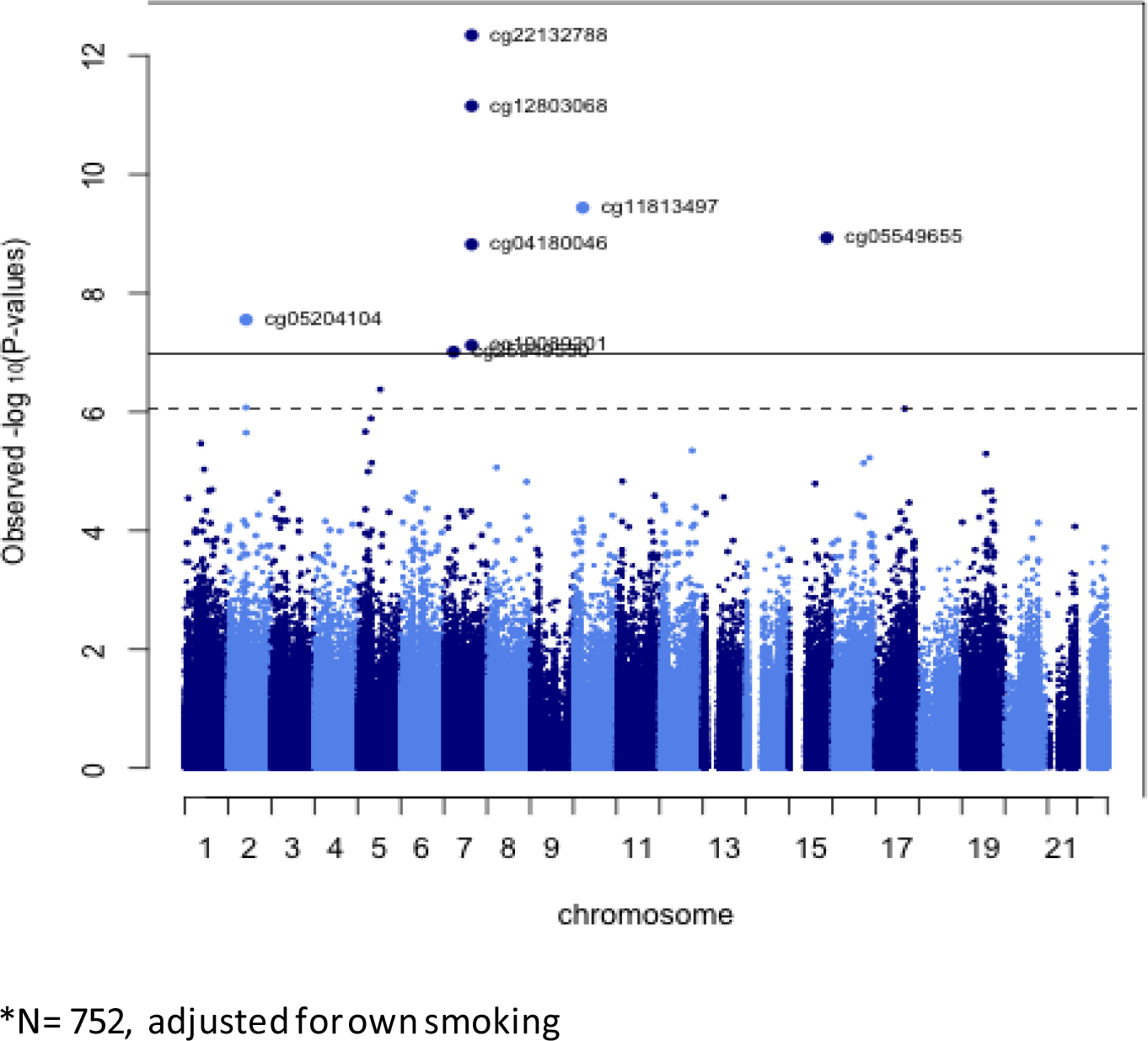
Manhattan plot for EWAS of prenatal smoke exposure in ALSPAC women*

Effects at the top CpGs surpassing FDR correction in the ALSPAC women were found to be consistent in direction although slightly attenuated in the follow up analysis approximately 18 years later (Pearson’s correlation coefficient, r = 0.92) (**Figure 2**, **Supplementary Figure 1**). Furthermore, there was remarkable consistency in the direction and magnitude of effects at these CpGs in blood samples of the ALSPAC men (r=0.77), with the exception of *DRD1* where effects were not as consistently replicated. We also compared for reference the effect of any maternal smoking on cord blood methylation in the ALSPAC birth cohort at these CpGs and, again, both magnitude and direction of effect was similar (r=0.92) with the exception of sites at *ARL4C* (cg05204104), *AHRR* (cg17924476) and *DRD4* (cg22807681). At these sites the difference in methylation was greater in the ALSPAC women than the newborns (**Figure 2**, **Supplementary Figure 1**).

**Figure 2:**
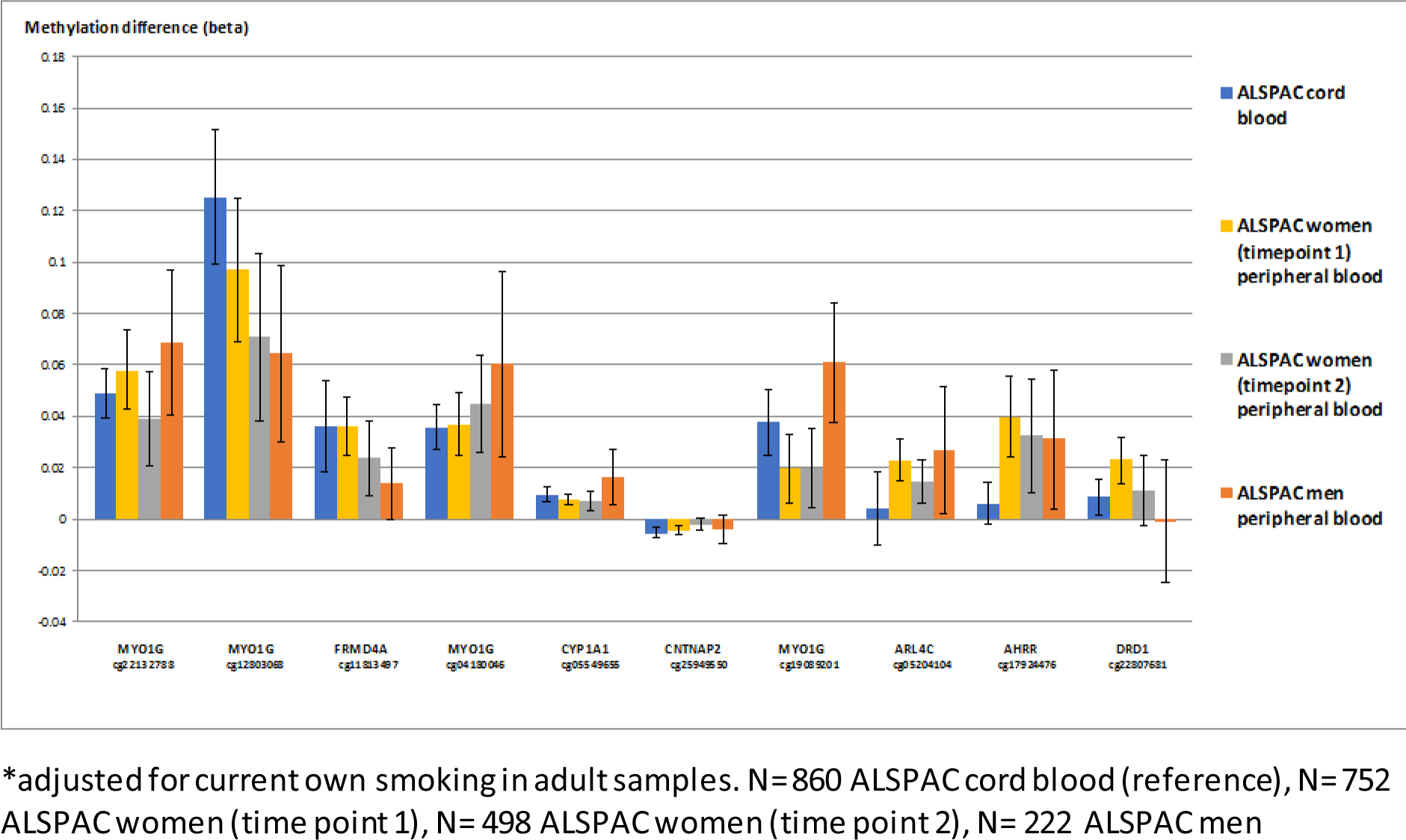
Replication of CpG sites observed below FDR (p<0.05) threshold in ALSPAC women at a later time point and in ALSPAC men*, and comparison with effect of prenatal smoking on cord blood methylation in ALSPAC children.

In addition, we found that among women with a mean age of 30 years (N=754), there was a strong signal of association above that expected by chance at CpG sites previously associated with prenatal smoke exposure in newborns (lambda = 2.62 vs. 1.14 for all CpG sites on the Illumina Infinium HM450 BeadChip; Wilcoxon rank sum test p-value < 2.2 × 10^−16^) (**Figure 3**, **Supplementary Figure 2**). Similarly, inflation of signals for prenatal smoke exposure was seen in these women 18 years later (lambda = 1.54, p-value = 5.6 × 10^−15^) and in the ALSPAC men (lambda = 1.19, p=2.2 × 10^−4^) (**Figure 3**), compared with all CpG sites on the HM450 BeadChip (**Supplementary Figure 2**).

**Figure 3:**
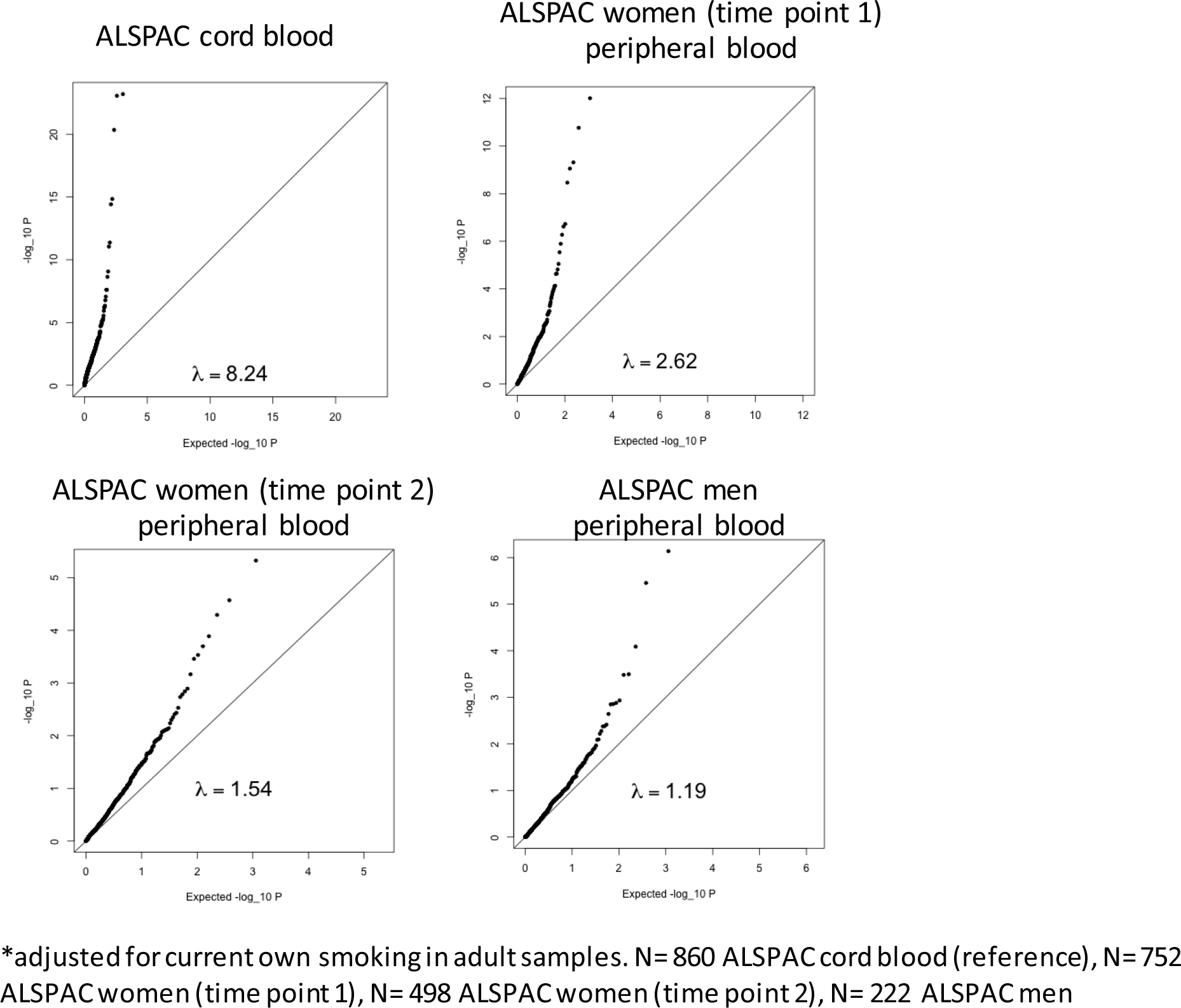
QQ plots and lambda values for 568 CpG sites previously found to be robustly associated with prenatal smoke in a cord blood meta-analysis in ALSPAC*

We found that a prenatal smoking methylation score, derived by combining methylation values at CpG sites previously associated with prenatal smoke exposure in cord blood of newborns, could predict whether the mothers of the ALSPAC women smoked during pregnancy with an AUC 0.69 (95% CI 0.67, 0.73). This was comparable with a score derived from CpG sites previously associated with prenatal smoking in peripheral blood of older children, which had an AUC 0.72 ((95% CI 0.69, 0.76; P for difference = 0.97) **(Figure 4)**.

**Figure 4:**
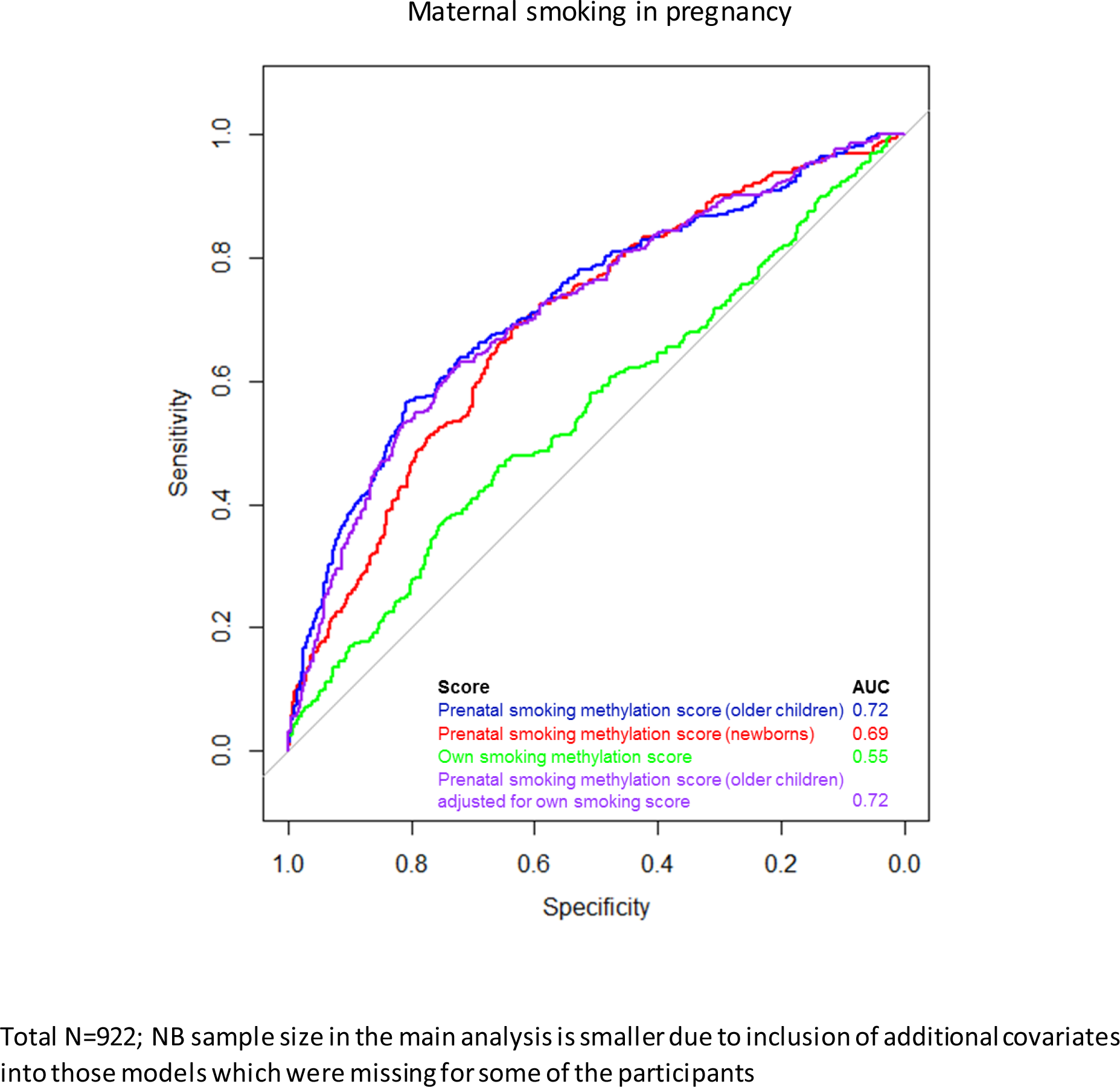
Receiver operating characteristic (ROC) curves of prenatal and own smoking methylation scores for discriminating maternal smoking in pregnancy

To determine the extent to which methylation associations with prenatal smoking were different from own smoking associations, we constructed a similar score derived from CpG sites previously associated with own smoking in adulthood. This score was only weakly associated with prenatal smoking (AUC 0.55, 95% CI 0.51, 0.59; P for difference = 2.1×10^−10^) **(Figure 4)**. In addition, the maternal smoking methylation score was able to predict with the same accuracy prenatal smoke exposure when adjusted for the offspring smoking methylation score (AUC 0.72 (95% CI 0.68, 0.75; P for difference = 0.43) **(Figure 4)**.

Furthermore, the prenatal smoking score derived from CpG sites associated with prenatal smoke exposure in older children was able to distinguish between individuals exposed to prenatal smoke regardless of whether or not they themselves smoked (p < 0.001; **Figure 5**). Interestingly, the prenatal smoking score derived from newborns by contrast was unable to distinguish between individuals exposed to prenatal smoke who did not themselves smoke and individuals unexposed to prenatal smoke who smoked themselves (p = 0.61).

**Figure 5:**
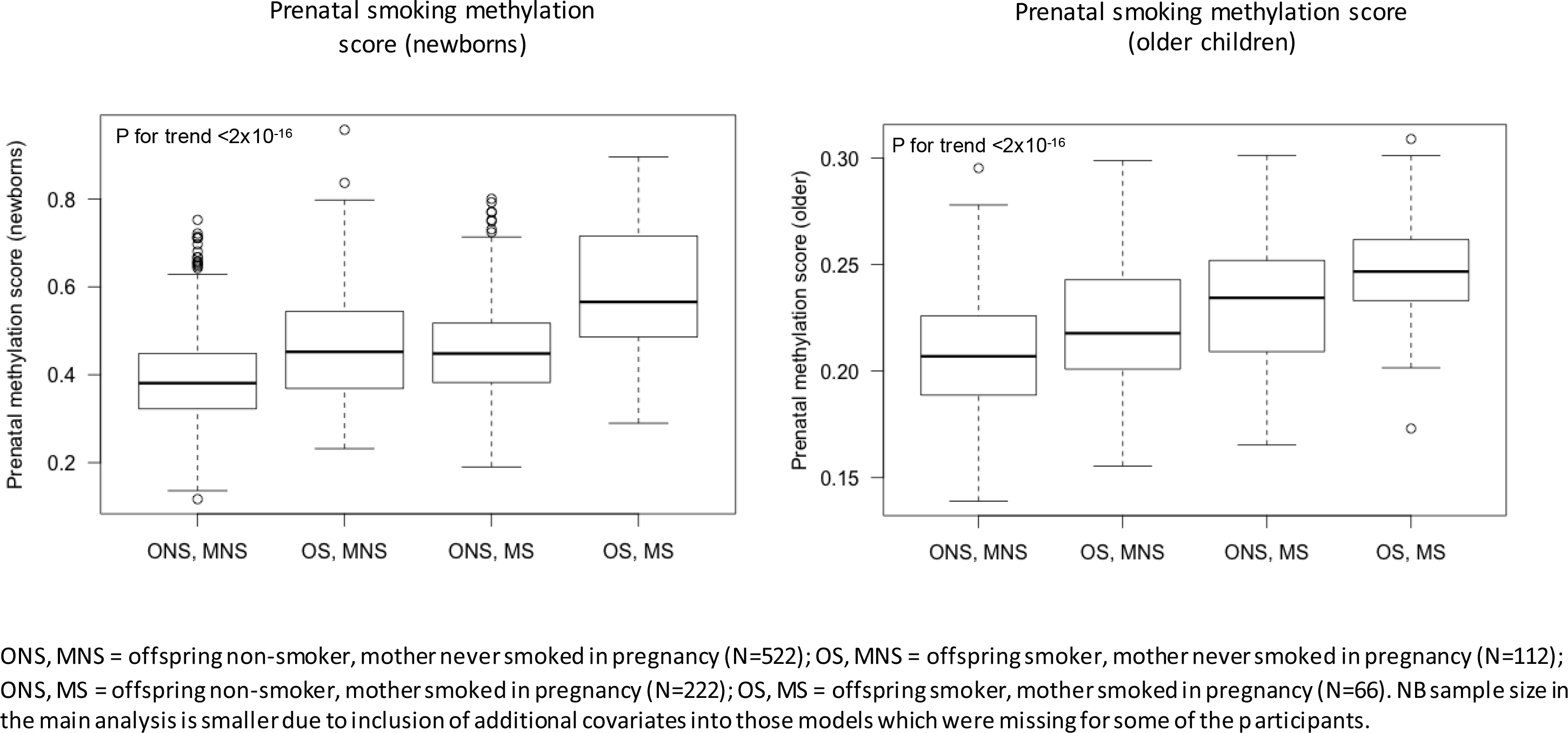
Box plots to assess differences in prenatal smoking methylation scores

As expected, the prenatal smoking methylation scores were higher in individuals (both smokers and non-smokers) exposed to prenatal smoking compared with non-smokers who were not exposed **(Figure 5)**. While the prenatal smoking methylation score derived from CpG sites previously associated with prenatal smoke exposure in cord blood of newborns (3) was not higher in non-smokers whose mothers smoked in pregnancy compared with smokers whose mothers didn’t smoke during pregnancy (difference in score = -0.07, p=0.61) **(Figure 5a)**, the score derived from CpG sites which were shown to persist in relation to prenatal smoke exposure based on an EWAS in older children (3) was higher in non-smokers whose mothers smoked in pregnancy compared with smokers whose mothers didn’t smoke in pregnancy (difference in score = 0.01, p=0.001) **(Figure 5b)**.

## Discussion

In a large longitudinal cohort with genome-wide methylation data, we identified 15 CpG sites that were differentially methylated in the peripheral blood of women over 30 years after exposure to prenatal smoking. Many of these CpG sites have been previously identified in relation to prenatal smoke exposure in the offspring at birth (3) and the majority showed long-term hypermethylation among the offspring of smokers.

Most of these signals remained in sensitivity analyses adjusted for own smoking and were observed in peripheral blood 18 years later (i.e. at around the age of 48 years). Furthermore, we replicated findings of a persistent methylation signal related to prenatal smoke exposure in peripheral blood among men in the ALSPAC cohort. For all these samples there was also a strong signal of association above that expected by chance at CpG sites previously associated with prenatal smoke exposure in newborns (3).

Furthermore, we found that a prenatal smoking score, derived by combining methylation values at these CpG sites, could adequately predict whether the mothers of the adults in ALSPAC smoked during pregnancy with an AUC 0.69 (95% CI 0.67, 0.73). A recent study identified a much stronger predictive ability of a prenatal smoking methylation score with AUC of 0.90 in a test set of cord blood obtained from newborns in the MoBa cohort (30). Furthermore, when we applied our prenatal smoking methylation score in the ARIES cord blood samples, it was shown to have an AUC of 0.89 in relation to maternal smoking. The difference in predictive ability is likely attributed to the 30-year difference in time since exposure and the generation of this methylation score using CpG sites identified in cord blood rather than adult peripheral blood. Furthermore, as the ARIES cord samples were included in the previous EWAS meta-analysis from which the prenatal smoking score was derived (3), this could produce some over-fitting of the estimates which would be less likely in the adult samples from ARIES which were not involved in this previous analysis.

We also derived a methylation score based on CpGs which showed evidence of a persistent difference in methylation in peripheral blood of offspring exposed to prenatal smoking (3) had a marginally higher AUC of 0.72 (95% CI 0.67, 0.73) and was also able to distinguish non-smokers whose mothers smoked in pregnancy from smokers whose mothers did not smoke during pregnancy.

Strengths of our study include the large sample size of women with reported maternal smoking in pregnancy for performing our initial EWAS analysis, the ability to adjust for own smoking status, the longitudinal assessment of differential methylation in a follow-up sample of these women, and the replication analysis in men from the same study.

Although there was evidence of persistence for methylation differences even after adjusting for own smoking status in the adult offspring, there are limitations to performing this type of adjustment analysis. As parental smoking is strongly associated with their offspring’s smoking initiation (31), own smoking serves as a possible mediator on the path between prenatal smoking and offspring DNA methylation. This method of adjusting for a potential mediator in standard regression models to estimate the direct effect of an exposure may produce spurious conclusions (32, 33). While an alternative method of using life course models previously provided more evidence for the hypothesis that maternal smoking in pregnancy is the ‘critical period’ for influencing persistent offspring methylation profiles (4), this method could not be applied here given the limited amount of information on maternal smoking reported by the ALSPAC men and women.

A further limitation relates to cell type heterogeneity, given that the ALSPAC samples were obtained from a variety of sources (white cells, whole blood and PBLs). To account for this, we incorporated surrogate variables into our models to account to adjust for technical batch and cell-type mixture in order to harmonize cellular variability of the samples (23).

In addition, information on prenatal smoke exposure in the ALSPAC men and women was recorded retrospectively by the adult offspring, rather than by prospective assessment and so may be subject to more misreporting. Furthermore, in the ALSPAC men and women, rates of maternal smoking in pregnancy were reported to be high in comparison with contemporary populations. This draws to question the relevance of identified associations. However, we have shown that many of the signals identified in adults were also present in cord blood of offspring measured prospectively in a more contemporary cohort with lower rates of maternal smoking in pregnancy (4).

The results of this study provide robust evidence that maternal smoking in pregnancy is associated with changes in DNA methylation that persist in the exposed offspring for many years after their prenatal exposure and across a number of tissues. Furthermore, these associations largely remain after adjusting for the previous smoking history of the adults themselves and are in accordance with earlier studies investigating prospectively assessed maternal smoking during pregnancy in relation to global DNA methylation levels (14, 15).

These findings could have useful applications in epidemiological studies, for example by using DNA methylation signatures as a refined exposure indicator and biosocial archive for historical exposure (5). Furthermore, persistent changes in DNA methylation might mediate at least some of the associations between smoke exposure in pregnancy and later life health outcomes (34). However, distinguishing mediation from other association-driving mechanisms (35) warrants further evaluation with the integration of analytical techniques such as two-step Mendelian randomization (36, 37), transcriptomic analysis and the profiling of target tissues (38).

## Acknowledgements

We are extremely grateful to all the families who took part in this study, the midwives for their help in recruiting them, and the whole ALSPAC team, which includes interviewers, computer and laboratory technicians, clerical workers, research scientists, volunteers, managers, receptionists, and nurses.

